# Design and validation of an exposure system for efficient inter-animal SARS-CoV-2 airborne transmission in Syrian hamsters

**DOI:** 10.1101/2022.11.18.517035

**Authors:** Philip J. Kuehl, Justin Dearing, Adam Werts, Jason Cox, Hammad Irshad, Edward G. Barrett, Sean N. Tucker, Stephanie N. Langel

## Abstract

SARS-CoV-2 is a highly transmissible respiratory pathogen whose main transmission route is airborne. Development of an animal model and exposure system that recapitulates airborne transmission of SARS-CoV-2 is integral for understanding the dynamics of SARS-CoV-2 spread in individuals and populations. Here we designed, built, and characterized a hamster transmission caging and exposure system that allows for efficient SARS-CoV-2 airborne transmission from an infected index animal to naïve recipients under unidirectional airflow, without contribution from fomite or direct contact transmission. To validate our system, we assessed a 1:1 or 1:4 ratio of infected index to naive recipient hamsters and compared their virological and clinical measurements after eight hours of airborne exposure. Airborne exposure concentrations and pulmonary deposited dose of SARS-CoV-2 in index and naïve hamsters, respectively, were similar in both groups. Daily nasal viral RNA levels, and terminal (day 5) lung viral RNA and infectious virus, and fecal viral RNA levels were statistically similar among 1:1 and 1:4 naive animals. However, virological measurements in the 1:4 naïve animals were more variable than the 1:1 naïve animals, likely due to hamster piling behavior creating uneven SARS-CoV-2 exposure during the grouped 1:4 airborne exposure. This resulted in slight, but not statistically significant, changes in daily body weights between the 1:1 and 1:4 naive groups. Our report describes a multi-chamber caging and exposure system that allowed for efficient SARS-CoV-2 airborne transmission in single and grouped hamsters. This system can be used to better define airborne transmission dynamics and test transmission-blocking therapeutic strategies against SARS-CoV-2.

**Importance:** The main route of SARS-CoV-2 transmission is airborne. However, there are few experimental systems that can assess airborne transmission dynamics of SARS-CoV-2 in vivo. Here, we designed, built, and characterized a hamster transmission caging and exposure system that allows for efficient SARS-CoV-2 airborne transmission in Syrian hamsters, without contributions from fomite or direct contact transmission. We successfully measured SARS-CoV-2 viral RNA in aerosols and demonstrated that SARS-CoV-2 is transmitted efficiently at either a 1:1 or 1:4 infected index to naïve recipient hamster ratio. This is meaningful as a 1:4 infected index to naïve hamster ratio would allow for simultaneous comparisons of various interventions in naïve animals to determine their susceptibility of infection by aerosol transmission of SARS-CoV-2. Our SARS-CoV-2 exposure system allows for testing viral airborne transmission dynamics and transmission-blocking therapeutic strategies against SARS-CoV-2 in Syrian hamsters.

## Introduction

Severe acute respiratory syndrome coronavirus 2 (SARS-CoV-2) is a highly transmissible respiratory pathogen spread by droplets and aerosols produced during coughing, sneezing, talking, and breathing (1–3). Airborne transmission of SARS-CoV-2 is dependent on multiple factors including duration and magnitude of viral shedding, force of exhalation, stability of the virus in aerosols, and immune status of infected individuals (4, 5). SARS-CoV-2 can spread from an infected individual to one naïve individual or to a group of naïve individuals, particularly in congregate residency settings where residents have long duration and close contact (2). Deciphering the dynamics of SARS-CoV-2 transmission is an important goal for the development of public health strategies and biological countermeasures to inhibit aerosol transmission.

Animal models of SARS-CoV-2 transmission have been integral in defining the impact of SARS-CoV-2 replication, shedding, and immunity on viral transmission. The most common animal used for SARS-CoV-2 transmission studies is the (Syrian) golden hamster. Golden hamsters are highly susceptible to SARS-CoV-2, recapitulate multiple aspects of disease, and can transmit SARS-CoV-2 efficiently by aerosols and fomites (6, 7). To study airborne transmission of SARS-CoV-2, we designed a chamber system where infected donor hamsters (index animals) are placed in an empty chamber (chamber 1) upstream of a chamber containing uninfected naïve hamsters (chamber 3). A connecting chamber (chamber 2) connects the index and recipient chambers without allowing physical or fomite contact between the hamsters while under constant unidirectional airflow controlled by vacuum from chamber 3. We aimed to validate our caging and exposure system by determining SARS-CoV-2 transmission dynamics from an infected index hamster to one naïve recipient hamster (1:1) or to a group of naïve recipient hamsters (1:4).

Here, we highlight our chamber system design that allows for efficient airborne transmission of SARS-CoV-2 in golden hamsters. Additionally, we report the virological responses and clinical outcomes of index animals and naïve recipient animals (in either a 1:1 or 1:4 index to naïve ratio). Our data demonstrate that SARS-CoV-2 is transmitted efficiently from infected index to naïve recipient animals regardless of the housing ratio using our caging and exposure system. While there are slight differences in viral replication kinetics, they do not result in significant differences between the naïve recipient treatment groups. Our data demonstrate that either ratio of index to naïve recipient golden hamsters can be used to study SARS-CoV-2 transmission dynamics. Developing animal models and exposure systems of SARS-CoV-2 transmission is key for future testing of therapeutic strategies that limit airborne viral transmission.

## Results

### Design and validation of a caging and exposure system for airborne transmission of SARS-CoV-2

Index hamsters were infected with 1×10^5^ TCID_50_ of SARS-CoV-2 by intranasal administration and housed individually. After 24 hours, infected index hamsters were placed in chamber 1 and either one or four naïve recipient hamsters were placed in chamber 3 (Figure 1A,B). Viral aerosol sampling was performed in the connector chamber (chamber 2) via glass microfiber filters (Figure 1A,B) at 1 L/min for a total of 8 hours. Total aerosol concentration of the filter samples determined by differential mass analysis resulted in an average concentration of 0.21 (±.08) μg/L_air_ for single (1:1) and 0.18 (±0.1) μg/L_air_ for grouped (1:4) hamsters (Figure 2A, Data set S1 in supplemental material). Additionally, particle size measurements were collected with a GRIMM aerosol spectrophotometer (Figure 1C). Regardless of the duration of measurement or the number of collections taken, the number of aerosol particles collected was insufficient for analysis of particle size. The low total aerosol concentration and failure to detect aerosol particle size likely reflects the level of filtration for the inlet air (HEPA filter) and the low mass concentration of aerosol generated by the index hamster.

**FIG 1.**
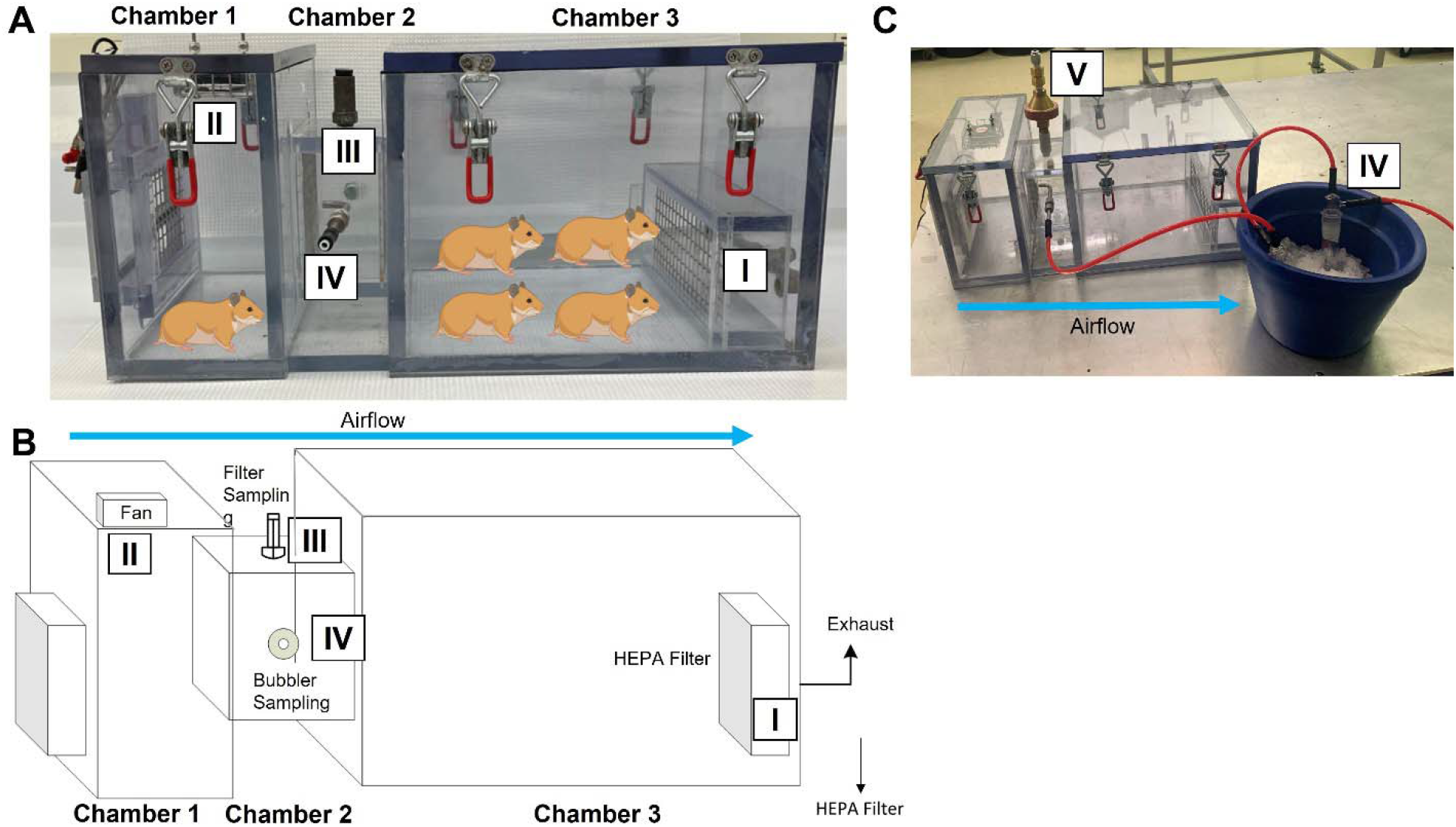
SARS-CoV-2 RNA detected in aerosols after infecting index hamsters using a three-chamber exposure system. (A, B) Chamber 1 housed the index hamsters and chamber 3 housed the naïve hamsters. A wire mesh screen was fit on either ends of the connector chamber (chamber 2) to separate the hamsters but allow for air passage. Unidirectional airflow was controlled by house exhaust flow (I) from chamber 3, drawing room air into chamber 1 through a HEPA filter. A recirculating fan (II) was placed on the top of chamber 1 to ensure homogeneity of airborne particles moving through chambers 2 and 3. Aerosols for total aerosol concentration testing were collected using glass microfiber filters located at the top of chamber 2 (III, ‘filter sampling’ in B). (A,B,C) Viral aerosol concentration was measured with an all glass impinger connected to a probe port on the side of chamber 2 (IV, ‘bubbler sampling’ in B). The particle size distribution was sampled with a GRIMM portable aerosol spectrometer from either III or IV that was placed on the top of chamber 2 (V).

**FIG 2.**
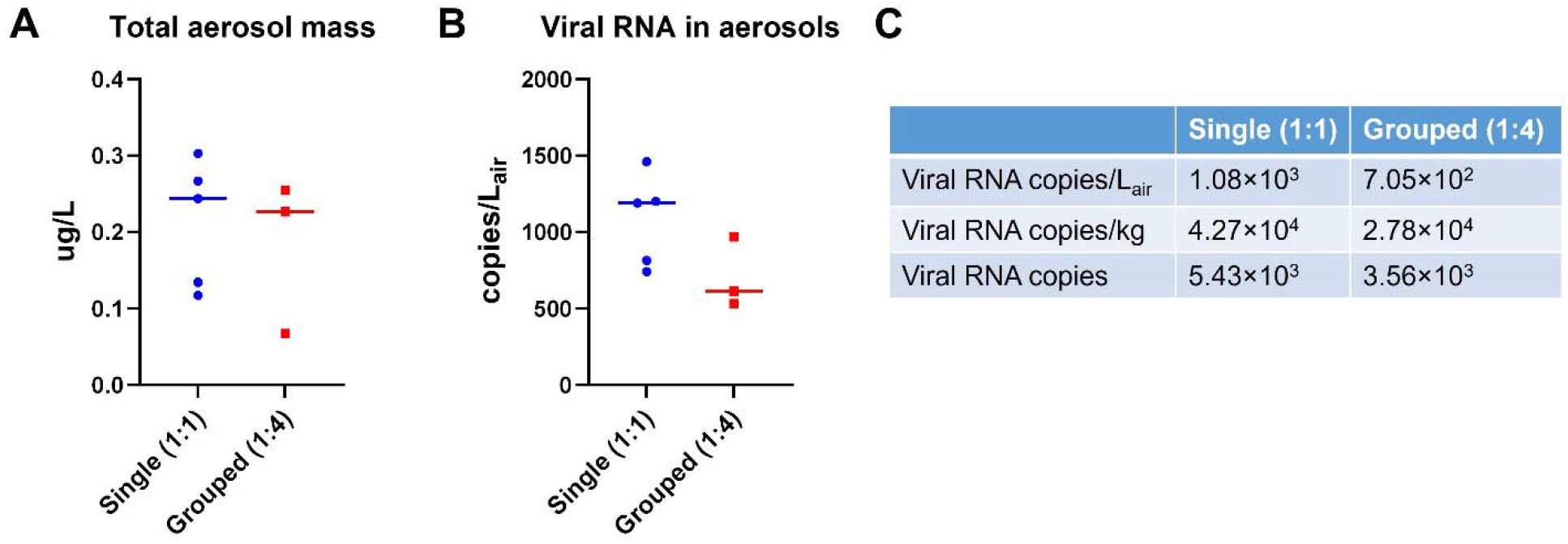
Total aerosol mass and viral RNA detected in aerosols in single (1:1) and group (1:4) hamsters. (A) Glass microfiber filters were used to collect aerosols and total aerosol mass was measured by differential mass analysis using an ultra-balance (0.001 mg sensitivity). (B) Viral RNA in aerosols were collected into an all glass impinger with Tris-EDTA buffer and measured via quantitative reverse transcription PCR (qRT-PCR). (C) Average SARS-CoV-2 viral RNA aerosols from single naïve (1:1) and group naïve (1:4) hamster chamber systems.

Aerosolized virus was collected in an all glass impinger (Figure 1A-C) filled with Tris-EDTA buffer downstream at a flow rate of 0.5 L/min for 8 hours. The analysis of the impinger material (genomic viral RNA copies) showed an average viral RNA aerosol concentration of 1.08×10^3^ (±2.29×10^2^) copies/L_air_ for single (1:1) and 7.05×10^2^ (±2.33×10^2^) copies/L_air_ for group (1:4) hamsters (Figure 2B, Data set S2 in supplemental material). Utilizing standard methods (8) and a 10% deposition fraction, we calculated an average pulmonary deposited dose of 3.71×10^4^ copies/kg body weight or 4.74×10^3^ copies per animal in all groups. Specifically, the 1:1 hamster group’s average aerosol concentration was 1.08×10^3^ copies/L_air_, 4.27×10^4^ copies/kg or 5.43×10^3^ copies and for the 1:4 animals 7.05×10^2^ copies/L_air_, 2.78×10^4^ copies/kg or 3.56×10^3^ copies (Figure 2C, Data set S3 in supplemental material). These data highlight the reproducibility of the aerosol concentration of SARS-CoV-2 viral RNA from the index animals.

### SARS-CoV-2-infected index hamsters transmit virus to single naïve (1:1) and group naïve (1:4) hamsters

Infected index hamsters had high levels of genomic (Figure 3A) and subgenomic (Figure 3B) viral RNA in nasal swabs at 1-day post-inoculation that decreased by day 5. However, for single (1:1) and group (1:4) naïve recipient hamsters, genomic and subgenomic viral RNA levels in nasal swabs increased at day 1 post exposure but did not reach comparable levels to index hamsters until day 3 post exposure, likely due to lower inoculation dose in these animals compared to index animals. A small proportion of the 1:4, but not 1:1, naïve recipient group hamsters had genomic and subgenomic viral RNA levels near the limit of quantification (LOQ) throughout the study demonstrating variability in SARS-CoV-2 transmission and replication in 1:4 naïve hamsters. Despite this, genomic and subgenomic viral RNA levels in nasal swabs were not significantly different between 1:1 and 1:4 naïve recipient hamsters at any time point. Overall, this demonstrates that SARS-CoV-2 was transmitted effectively from infected index hamsters to single naïve and group naïve recipient hamsters after 8 hours of airborne exposure.

**FIG 3.**
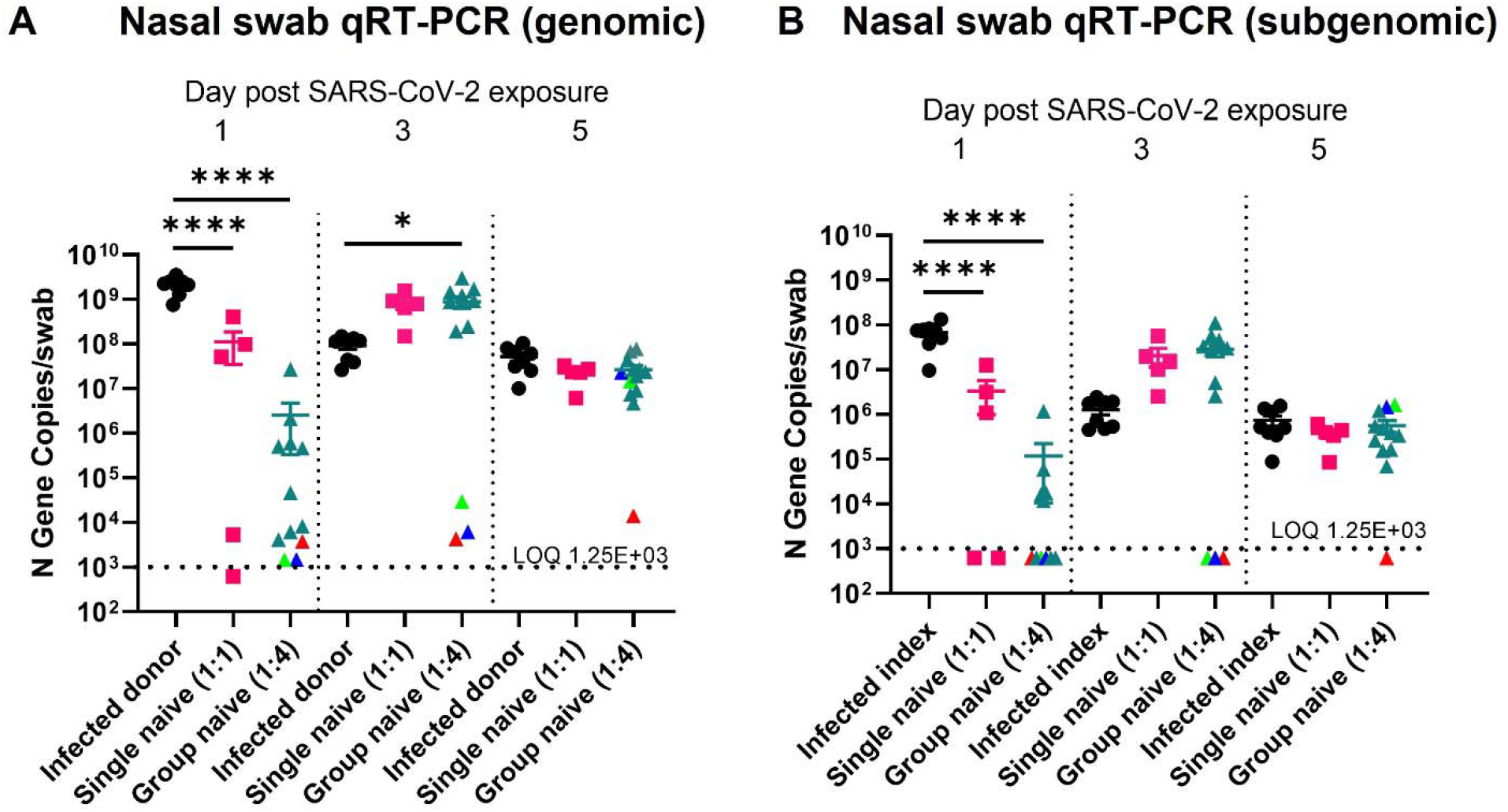
SARS-CoV-2 transmission and replication kinetics in infected index, single (1:1) naive, and group (1:4) naïve recipient hamsters. Nasal swabs were collected on days 1, 3, and 5 postinoculation in infected index hamsters and on days 1, 3, and 5 in naïve recipient hamsters after exposure to infected index hamsters in airborne transmission chambers. Viral genomic (A) and subgenomic (B) RNA levels were determined by quantitative reverse transcription PCR (qRT-PCR) of the nucleocapsid (N) gene. The different color triangles (red, green, blue) for the group naïve (1:4) data points represent animals who remained near the limit of quantification (LOQ) at more than one time point (each animal retains the same color for each day post SARS-CoV-2 exposure). Daily qRT-PCR data were analyzed by a one-way ANOVA using Tukey’s multiple comparisons. Error bars represent the standard error of the mean (SEM). *P < 0.05, ****P< 0.0001.

### Levels of lung viral RNA and infectious virus are more variable in group (1:4) naïve recipient hamsters

Lung genomic and subgenomic viral RNA (Figure 4A) and infectious virus (Figure 4B) were high in all groups and not significantly different between infected index, single naïve (1:1), and group naïve (1:4) hamsters at necropsy (day 5). Fecal viral genomic RNA levels (Figure 4C), collected at necropsy (day 5) were also assessed and were not significantly different between infected index and naïve recipient groups. However, a greater proportion of the 1:4 naïve hamsters had lung viral RNA and infectious virus levels near the LOQ compared to 1:1 naïve hamsters, demonstrating variable transmission efficiency and/or virus replication post exposure to index hamsters in the 1:4 naïve hamsters. Interestingly, the two animals with lung subgenomic RNA and infectious viral titers near the LOQ did not have fecal viral genomic RNA near the LOQ. This suggests that in hamsters, like humans (9), viral RNA or infectious virus in the respiratory tract does not predict viral RNA levels in feces.

**FIG 4.**
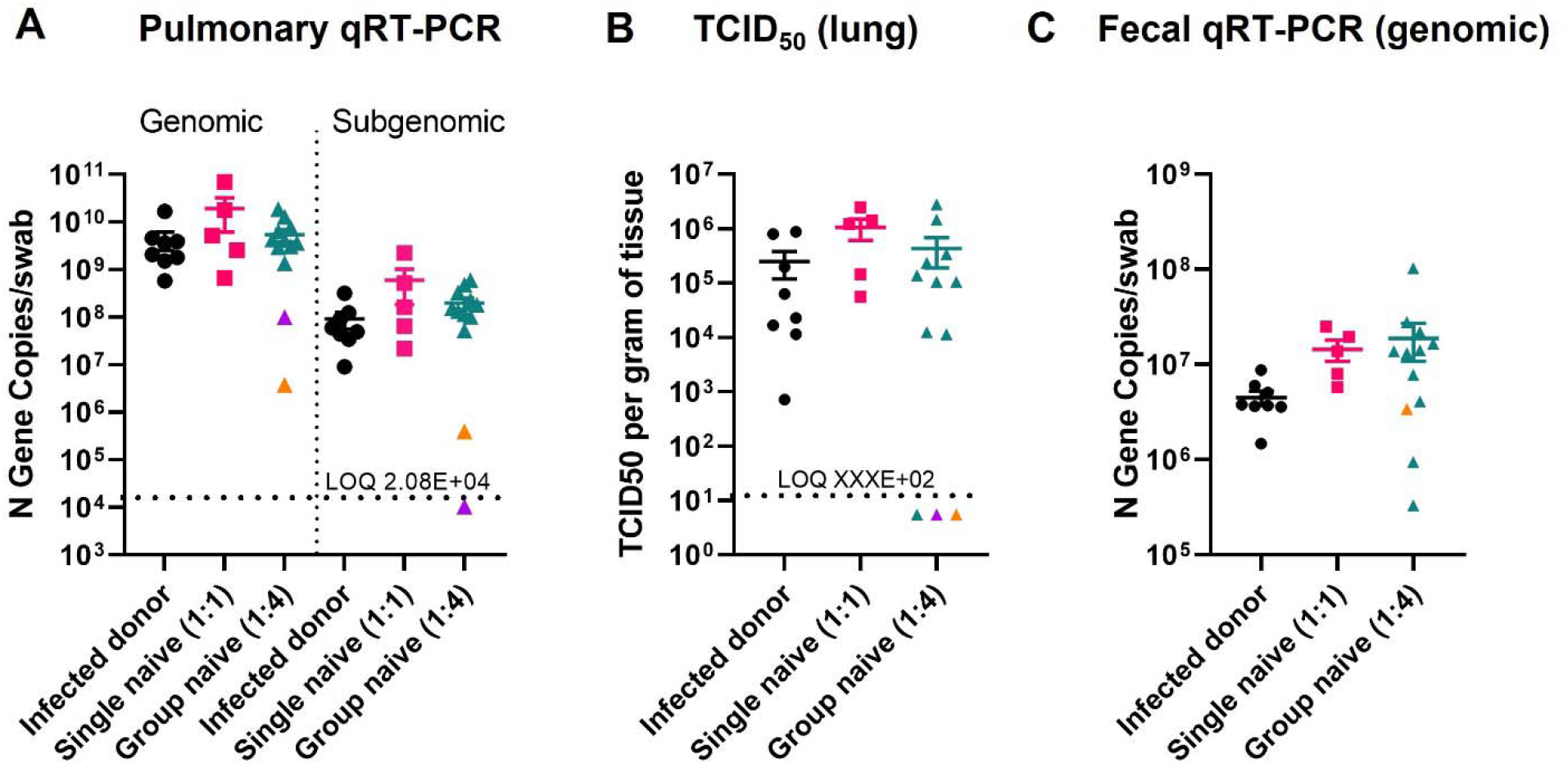
SARS-CoV-2 viral RNA levels in lung and feces between index, single (1:1) naive, and group (1:4) naïve recipient hamsters at necropsy (day 5). Lung tissue (A) and feces (C) were collected at necropsy (day 5) and genomic (lung and feces) and subgenomic (feces only) RNA was isolated for SARS-CoV-2 detection by quantitative reverse transcription PCR (qRT-PCR) of the nucleocapsid (N) gene. Infectious viral titers in lung tissue (B) were determined by TCID_50_. The dotted line represents the limit of quantification (LOQ). The different color triangles for the group naïve (1:4) data points represent animals who had lung subgenomic RNA and infectious viral titers near the LOQ (each animal retains the same color for each day post SARS-CoV-2 exposure). Data were analyzed by a one-way ANOVA using Tukey’s multiple comparisons.

### Body weights and lung weights were similar between single (1:1) and group (1:4) naïve recipient hamsters

Body weights as a percent of pre-SARS-CoV-2 exposure day 0 were measured in all groups. Infected index animals lost significantly more weight over the course of the experiment and had significantly lower terminal body weights (Figure 5A,B). Body weights over time and at necropsy were not significantly different between single (1:1) and group (1:4) naïve recipient hamsters.

**FIG 5.**
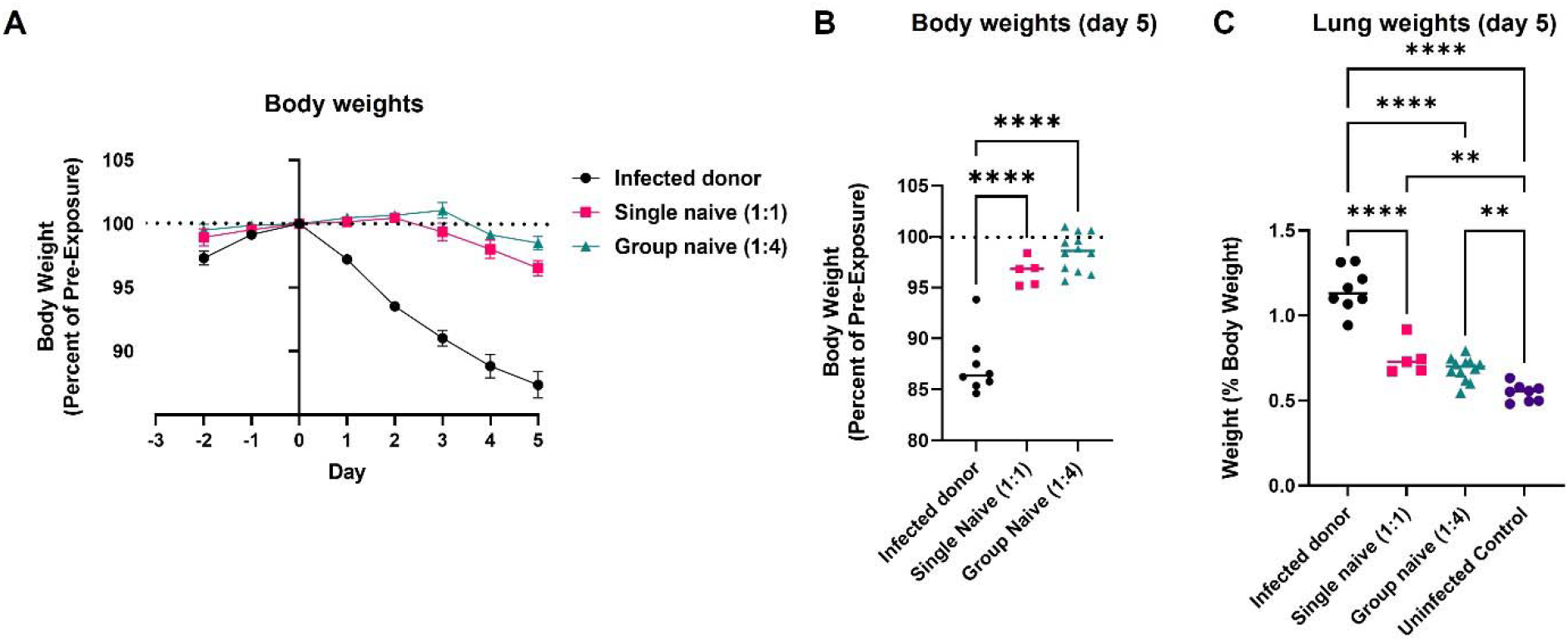
Clinical outcomes were similar between single (1:1) and group (1:1) naïve recipient hamsters. Daily weight changes (A) and terminal body weights (B) were determined by the percent of day 0 weight (relative to SARS-CoV-2 exposure). Terminal lung weights (C) were determined at necropsy. In B and C data were analyzed by a one-way ANOVA using Tukey’s multiple comparisons, respectively. Error bars in A represent the standard error of the mean (SEM). **P < 0.01, ****P< 0.0001.

Body weight-normalized lung weights, an indirect measure of pulmonary inflammation, were compared amongst all groups including uninfected control hamsters (Figure 5C). All infected groups had significantly higher lung weights compared to uninfected control hamsters. However, lung weights were greatest in the infected index hamsters. There were no significant differences in lung weights between 1:1 or 1:4 naïve hamsters.

## Discussion

SARS-CoV-2 transmission dynamics play a significant role in the ongoing COVID-19 pandemic. Animal models of SARS-CoV-2 transmission allow for the study of environmental and host factors that impact viral transmission and replication, as well as testing of therapeutic strategies (10). However, viral transmission characteristics differ for aerosols compared to direct contact or fomite exposure (6). Therefore, designing caging and exposure systems that allow for airborne transmission of SARS-CoV-2 in animals without contributions from other routes of exposure is important to understand transmission dynamics and test therapeutic strategies that inhibit airborne transmission. Here, we designed and characterized a caging and exposure system for SARS-CoV-2 airborne transmission and validated it using infected index hamsters cohoused, but physically separated from either one (single) or four (grouped) naïve recipient hamsters.

Previous groups have demonstrated that SARS-CoV-2 is transmitted efficiently from infected index to naïve recipient hamsters. Sia and colleagues infected index hamsters with 8×10^4^ TCID_50_ of SARS-CoV-2 and after 24 hours placed each index animal in a new cage cohoused with one naïve hamster (7). SARS-CoV-2 was detected from cohoused hamsters at 1-day post cohousing and reached peak viral loads in nasal washes at day 3. However, this experiment could not distinguish between airborne and fomite transmission considering naïve hamsters were cohoused in a way that allowed contact with infected index hamsters. In a more recent study, Port and colleagues designed a chamber system where SARS-CoV-2 transmission was observed under constant airflow of airborne particles <5μm over a 200 cm distance (11). Indeed, SARS-CoV-2 transmission could occur at this distance between Syrian hamsters even within 1 hour at a 2:2 ratio (2 infected index and 2 recipient naïve hamsters). Investigators were also able to demonstrate rates of airborne transmission were different between SARS-CoV-2 variants (11). However, SARS-CoV-2 RNA levels in aerosols were not quantified.

We collected aerosols using glass impingers and quantified SARS-CoV-2 viral RNA. We observed similar amounts of SARS-CoV-2 viral RNA in aerosols between chambers of the 1:1 and 1:4 treatment groups. This demonstrated that index animals inoculated with a consistent dose of SARS-CoV-2 (1×10^5^ TCID_50_/animal) transmitted similar amounts of SARS-CoV-2 in aerosols to naïve hamsters in our unidirectional airflow exposure system. At day 1 post exposure, nearly all the naïve hamsters in 1:1 and 1:4 groups had viral RNA levels above the LOQ. Interestingly, in the 1:4 naïve recipient group, there were hamsters on day 3 (3 hamsters) and 5 (1 hamster) that had nasal swab viral RNA levels that remained near the LOQ. This demonstrates increased variability of SARS-CoV-2 exposure and/or replication in the 1:4 naïve compared to the 1:1 naive recipient hamsters. Hamsters are known to group together when cohoused and were observed to be sleeping in piles in some transmission chambers during the exposure period. It is possible that the one hamster in the 1:4 naïve recipient group with consistently low levels of viral RNA was piled underneath the other hamsters during cohousing with the infected index hamsters, and therefore did not receive a similar inoculum dose via aerosols as its cage mates. Despite this, statistically there were no differences in nasal swab viral RNA levels between 1:1 or 1:4 naïve treatment groups.

Like virological measurements in nasal swabs, lung viral RNA levels, lung infectious viral titers, and fecal viral RNA levels were more variable in the 1:4 compared to the 1:1 naïve recipient group. The explanation is expected to be similar as the nasal swabs; hamsters piling together may lead to uneven distribution of transmitted SARS-CoV-2 during airborne exposure from infected index hamsters. Additionally, the mean body weights (as a percent of pre-exposure) were numerically higher in 1:4 compared to 1:1 naïve recipient. This suggests that the potential lower SARS-CoV-2 exposure in some of the 1:4 hamsters resulted in less body weight loss over the course of the experiment. However, for both virological measurements and clinical signs, the overall levels were not significantly different between the 1:1 and 1:4 naïve recipient groups. Therefore, it is unlikely that the variability level observed in the 1:4 naïve recipient groups will significantly impact a study’s design and/or outcomes, depending on the research question.

Developing animal models and exposure systems of SARS-CoV-2 transmission is integral to define transmission kinetics and develop therapeutics that either prevent or stop viral transmission to naïve individuals. Here, we designed a caging and exposure system and demonstrated that SARS-CoV-2 is transmitted efficiently at either a 1:1 or 1:4 infected index to naïve recipient hamster ratio. From a practical standpoint, using a 1:1 ratio would require a large number of chambers, and might not be feasible for comparing interventions effectively. A 1:4 ratio would allow simultaneous comparisons of various interventions in naïve animals to determine their susceptibility of infection by inter-animal aerosol transmission of virus. In conclusion, we designed, built, and validated an exposure system that can be used to better define viral airborne transmission dynamics and to test transmission-blocking therapeutic strategies against SARS-CoV-2.

## Materials & Methods

### Characteristics of the transmission chamber system

The exposure system allowed unidirectional air flow from SARS-CoV-2 infected index hamsters in the first chamber (4”x10”x9”) to naïve recipient hamsters in the third chamber (10”x10”x9”). A second chamber (4”x5”x5”) connected chambers 1 and 3 and was raised 1 inch off the ground to prevent feces and fomite transfer. The chambers were made of polycarbonate. Wire mesh screens (0.25”x 0.25” holes) were fit on either ends of the connector chamber to separate the hamsters but allow for air passage. The first chamber was fit with a recirculating fan to ensure homogeneity of airborne particles moving through chambers 2 and 3. The fan had a wire cage around it to prevent animal access. Unidirectional flow was controlled by house exhaust flow from the chamber 3 at 5 L/minute, drawing room air into chamber 1 through a HEPA filter.

Sampling was performed in the connection chamber via probe ports. Filter samples (glass microfiber filters, Whatman Type GF/A) located at the port on top of chamber 2 were collected at 1 L/min to determine the total aerosol concentration. Filters were collected for between 1 and 8 hours to collect sufficient material for differential mass analysis via ultra-balance (0.001 mg sensitivity). Particle size distribution was sampled with a GRIMM portable aerosol spectrometer (Model 1.109, GRIMM Technologies, Inc. Germany). Viral aerosol concentration was measured with an all glass impinger (Midget Impinger, ACE Glass Inc) located at the port on the side of chamber 2. Impingers were filled with 10 mL of TE Buffer and collected at a flow rate of 0.5 L/min for up to 8 hours. Remaining TE buffer was weighed (to normalize for evaporation during collection) and assayed for SARS-CoV-2 by qRT-PCR.

### Animals

Male Syrian hamsters (*Mesocricetus auratus*) 12 to 14 weeks of age (weight range 106 to 136 g) were sourced from Charles River Laboratory. Animal work was performed at Lovelace Biomedical, with approval from the Institutional Animal Care and Use Committee (IACUC FY20-117E-E5). Hamsters were singly housed in filter-topped cage systems and were supplied with a certified diet, filtered municipal water, and dietary and environmental enrichment prior to transmission experiments.

Index animals were anesthetized by an intraperitoneal (IP) injection of a 10 mg/kg ketamine and 5 mg/kg xylazine cocktail for inoculation. Vaporized inhalant isoflurane gas (maximum of 5%) was used if additional restraint was necessary. Upon reaching the desired plane of anesthesia, the animal was held upright and intranasally inoculated dropwise with 100 μL per naris of a 5×10^5^ TCID_50_/mL SARS-CoV-2 in Dulbecco’s Modified Eagle Medium (DMEM) inoculum for a total of approximately 1×10^5^ TCID_50_/animal. All infected animals were monitored cage side until their righting reflex was restored.

Approximately 24 hours later, the transmission chambers were prepared, and animals were loaded. During the 8-hour transmission experiment, bedding and enrichment were removed however a DietGel® Boost (Clear H_2_O, product code: 72-04-5022) was provided ad libitum in both chambers 1 and 3 to meet the food and water needs of the animals throughout their 8-hour duration in the chamber. Naïve recipient hamsters were loaded and secured in chamber III before the infected index animals were handled. Animals were observed regularly while on exposure to monitor welfare. At the end of the 8-hour exposure, naïve hamsters were returned to their home cages before the infected index animals were handled. Each time animals were handled, technicians put on new personal protective equipment to reduce risk of SARS-CoV-2 transfer from handling infected index animals.

Nasal swabs were collected with Maxapplicators (Plasdent) with 0.5mm ultra fine fiber head tips. The animal was briefly anesthetized with vaporized inhalant isoflurane anesthetic and the swab was inserted 2 to 4mm into either the right or left naris and circled/twisted gently. The swab was then removed, placed into a Safe-Lock Eppendorf tube containing 1 mL of Trizol (TRI reagent, Millipore Sigma), and cut at the hinge point. Swabs were flash frozen at −80°±10°C for future RNA isolation. The nares sampled were alternated at each timepoint.

Animals were euthanized via intraperitoneal injection with a barbiturate overdose of pentobarbital (390 mg/mL) diluted in 1:10 in normal saline. After confirmation of death, the lungs were removed, weighed, and placed into a Safe-Lock Eppendorf tube and flash frozen at −80°±10°C for future RNA isolation and virus quantification by TCID_50_. Additionally, 2-3 fecal pellets were collected from the rectum, weighed, and placed into a Safe-Lock Eppendorf tube and flash frozen at −80°±10°C for future RNA isolation.

### RNA isolation of nasal swab, lung samples and feces

Lung tissue and feces samples were weighted (≤75 mg) and homogenized with beads using a Tissue Lyzer (Qiagen) in 1 ml of TRI reagent before RNA was isolated and purified from tissue samples using the Direct-Zol 96-RNA kit (Zyma Research). Nasal swabs were removed from −80°±10°C storage and mixed with TRI reagent (between 0.3 and 1 mL). Samples were mixed by vortexing briefly and allowed to incubate at room temperature for 15 minutes prior to RNA isolation.

### Detection of SARS-CoV-2 genomic and subgenomic RNA by qRT-PCR

Copies of the SARS-CoV-2 N gene (genomic) were measured by qRT-PCR TaqMan Fast Virus 1-step assay (Applied Biosystems). SARS-CoV-2 specific primers and probes from the 2019-nCoV RUO Assay kit (Integrated DNA Technologies) were used: (L Primer: TTACAAACATTGGCCGCAAA; R primer: GCGCGACATTCCGAAGAA; probe:6FAM-ACAATTTGCCCCCAGCGCTTCAG-BHQ-1). Reactions were carried out on a Stratagene MX3005P or BioRad CFX384 Touch instrument according to the manufacturer’s specifications. A semi-logarithmic standard curve of synthesized SARS-CoV-2 N gene RNA (LBRI) was obtained by plotting the Ct values against the logarithm of cDNA concentration and used to calculate SARS-CoV-2 N gene in copies per gram of tissue.

Copies of SARS-CoV-2 E gene (subgenomic) were measured by qRT-PCR with TaqMan Fast Virus 1-step assay (Thermo Fisher). SARS-CoV-2 specific primers and probes sgLeadSARSCoV2-F 5’-CGATCTCTTGTAGATCTGTTCTC-3’, E Sarbeco R: 5’-ATATTGCAGCAGTACGCACACA-3’, E_Sarbeco_P1: 6FAM-ACACTAGCCATCCTTACTGCGCTTCG-BHQ-1, E Sarbeco F: 5’-ACAGGTACGTTAATAGTTAATAGCGT-3’. Reactions were carried out on a Stratagene MX3005P or BioRad CFX384 Touch instrument according to the manufacturer’s specifications.

A semi-logarithmic standard curve of synthesized SARS-CoV-2 E gene RNA (LBRI) was obtained by plotting the Ct values against the logarithm of cDNA concentration and used to calculate SARS-CoV-2 N gene in copies per gram of tissue.

### Statistical analysis and aerosol concentration calculations

The methods used for determining significance were a one-way ANOVA and Tukey’s multiple comparisons using GraphPad Prism Software.

Aerosol dose for the naïve hamsters was calculated using an inhaled dose equation (8):

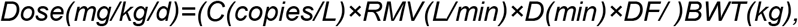

where C is the concentration (copies/L) in air inhaled (aerosol concentration), RMV, respiratory minute volume (L/min); D, duration of exposure (minutes); DF, deposition fraction; BWT, body weight (kg). For example, the aerosol concentration was the average aerosol concentration (copies/L_air_), the exposure duration (480 minutes), average group body weight (1:1 and 1:4 groups) and the assumed deposition fraction of 10%. While there are no specific data for a 10% deposition fraction for hamsters it is likely that the deposition will be like those used for rats and mice.

Particle size distribution samples all showed insufficient particle collections (background levels) across the entire particle size range for the GRIMM instrument. Therefore, no particle size reduction / analysis was performed.

## Author contributions

A.W., E.G.B., S.N.T, and S.N.L. conceptualized and designed the animal experiments. P.J.K., A.W., J.C., and H.I. conceptualized and designed the airborne transmission chamber. A.W. J.D., and J.C. carried out the transmission experiments. P.J.K., A.W., and S.N.L analyzed the data. P.J.K. and S.N.L. wrote the original draft and all authors reviewed and edited the draft.

## REFERENCES

1. Greenhalgh T, Jimenez JL, Prather KA, Tufekci Z, Fisman D, Schooley R. 2021. Ten scientific reasons in support of airborne transmission of SARS-CoV-2. Lancet 397:1603–1605.

2. Centers for Disease Control and Prevention. Scientific Brief: SARS-CoV-2 Transmission. 2021. https://www.cdc.gov/coronavirus/2019-ncov/science/science-briefs/sars-cov-2-transmission.html.

3. Wang CC, Prather KA, Sznitman J, Jimenez JL, Lakdawala SS, Tufekci Z, Marr LC. 2021. Airborne transmission of respiratory viruses. Science 373:eabd9149.

4. Lee EC, Wada NI, Grabowski MK, Gurley ES, Lessler J. 2020. The engines of SARS-CoV-2 spread. Science 370:406–407.

5. Meyerowitz EA, Richterman A, Gandhi RT, Sax PE. 2021. Transmission of SARS-CoV-2: A Review of Viral, Host, and Environmental Factors. Ann Intern Med 174:69–79.

6. Port JR, Yinda CK, Owusu IO, Holbrook M, Fischer R, Bushmaker T, Avanzato VA, Schulz JE, Martens C, van Doremalen N, Clancy CS, Munster VJ. 2021. SARS-CoV-2 disease severity and transmission efficiency is increased for airborne compared to fomite exposure in Syrian hamsters. Nat Commun 12:4985.

7. Sia SF, Yan LM, Chin AWH, Fung K, Choy KT, Wong AYL, Kaewpreedee P, Perera R, Poon LLM, Nicholls JM, Peiris M, Yen HL. 2020. Pathogenesis and transmission of SARS-CoV-2 in golden hamsters. Nature 583:834–838.

8. Tepper JS, Kuehl PJ, Cracknell S, Nikula KJ, Pei L, Blanchard JD. 2016. Symposium Summary: “Breathe In, Breathe Out, Its Easy: What You Need to Know About Developing Inhaled Drugs”. Int J Toxicol 35:376–92.

9. Natarajan A, Zlitni S, Brooks EF, Vance SE, Dahlen A, Hedlin H, Park RM, Han A, Schmidtke DT, Verma R, Jacobson KB, Parsonnet J, Bonilla HF, Singh U, Pinsky BA, Andrews JR, Jagannathan P, Bhatt AS. 2022. Gastrointestinal symptoms and fecal shedding of SARS-CoV-2 RNA suggest prolonged gastrointestinal infection. Med (N Y) 3:371–387.e9.

10. de Vries RD, Rockx B, Haagmans BL, Herfst S, Koopmans MP, de Swart RL. 2021. Animal models of SARS-CoV-2 transmission. Curr Opin Virol 50:8–16.

11. Port JR, Yinda CK, Avanzato VA, Schulz JE, Holbrook MG, van Doremalen N, Shaia C, Fischer RJ, Munster VJ. 2022. Increased small particle aerosol transmission of B.1.1.7 compared with SARS-CoV-2 lineage A in vivo. Nat Microbiol 7:213–223.

